# Formal Recognition and Classification of Gene Transfer Agents as Viriforms

**DOI:** 10.1101/2022.06.10.494566

**Authors:** Roman Kogay, Sonja Koppenhöfer, J. Thomas Beatty, Jens H. Kuhn, Andrew S. Lang, Olga Zhaxybayeva

## Abstract

Morphological and genetic features strongly suggest that gene transfer agents (GTAs) are caudoviricete-derived entities that have evolved in concert with cellular genomes to such a degree that they should not be considered viruses. Indeed, GTA particles resemble caudoviricete virions but, in contrast to caudoviricetes (or any viruses), GTAs can encapsidate at best only part of their own genomes, are induced solely in small subpopulations of prokaryotic host cells and are transmitted vertically as part of cellular genomes during replication and division. Therefore, the lifecycles of GTAs are analogous to virus-derived entities found in parasitoid wasps, which have recently been recognized as non-virus entities and therefore reclassified as viriforms. We evaluated three distinct, independently exapted GTA groups for which the genetic basis for GTA particle production has been established. Based on the evidence, we outline a classification scheme for these viriforms.

## INTRODUCTION

In 2021, the International Committee on Taxonomy of Viruses (ICTV) ratified a taxonomic proposal to formally accept a new operational definition of the term “virus” (Koonin et al. 2021; Kuhn et al. 2020; Walker et al. 2021). Consequently, the most current version of the International Code of Virus Classification and Nomenclature (ICVCN) states that viruses are

> “ … a type of MGEs [mobile genetic elements] that encode at least one protein that is a major component of the virion encasing the nucleic acid of the respective MGE and therefore the gene encoding the major virion protein itself; or MGEs that are clearly demonstrable to be members of a line of evolutionary descent of such major virion protein-encoding entities” (ICVCN Rule 3.3) (International Committee on Taxonomy of Viruses 2022; Kuhn et al. 2020).

This definition also formalized the postulate that some MGEs, long understood by the general virology community to be distinct from viruses, are indeed distinct. At the time, the ICTV had already classified viroids and satellite nucleic acids in taxa separate from viral taxa (in families/genera with names that end with suffixes -*viroidae*/-*viroid* and -*satellitidae*/-*satellite*, respectively, as opposed to -*viridae*/-*virus*) (International Committee on Taxonomy of Viruses 2022), and these elements were logically placed into the perivirosphere rather than the orthovirosphere (Koonin et al. 2021; Kuhn et al. 2020).

The adoption of the new virus definition brought into question the taxonomic standing of one official virus family, *Polydnaviridae*. Indeed, entities classified into this polyphyletic family fundamentally deviate from MGEs fulfilling the virus definition because “polydna” particles encapsidate multiple segments of circular double-stranded DNAs that, however, do not encode the entire “polydna” genomes. Instead, the genomes are permanently endogenized into the “polydna” host (i.e., parasitoid wasp) genomes and inherited vertically. The resultant non-mobile nonviral entities are used by the wasps to deliver immunomodulatory genes into insects that serve as prey for the wasps (Drezen et al. 2017; Herniou et al. 2013). “Polydna” entities are likely evolutionarily derived from various groups of insect viruses, including nudivirids (Darboux et al. 2019; Drezen et al. 2017; Gauthier et al. 2018; Petersen et al. 2022; Strand and Burke 2020; Thézé et al. 2011), but, because they have lost the ability to replicate and instead have been fully exapted by their wasp hosts, they have left the virosphere altogether (Koonin and Krupovic 2018; Koonin et al. 2021). Consequently, in 2021, the ICTV recognized “polydna” entities as representatives of a new MGE category distinct from viruses called “viriforms” (Koonin et al. 2021; Kuhn et al. 2020; Walker et al. 2021), and reclassified *Polydnaviridae* as (still polyphyletic) *Polydnaviriformidae* (Kuhn et al. 2021; Walker et al. 2022). In the ICVCN, viriforms are defined operationally as

> “ … a type of virus-derived MGEs that have been exapted by their organismal (cellular) hosts to fulfill functions important for the host life cycle; or MGEs that are derived from such entities in the course of evolution” (ICVCN Rule 3.3) (International Committee on Taxonomy of Viruses 2022; Kuhn et al. 2020).

Importantly, the following comment was added to the Rule 3.3:

“Gene transfer agents (GTAs) and the MGEs previously classified in the family *Polydnaviridae* are considered to be viriforms in classification and nomenclature” (International Committee on Taxonomy of Viruses 2022; Kuhn et al. 2020).

Notably, there are no discernible evolutionary relationships between GTAs and polydnaviriformids. The term “viriform”, similar to the term “virus”, is an umbrella term for certain MGEs with comparable lifecycles and properties; it is currently applied to six realms of MGEs that are not evolutionary related to each other.

Based on the properties of entities referred to as “GTAs” in the literature (reviewed in (Lang et al. 2012; Lang et al. 2017) we define GTAs as viriforms with the following features:

1. GTAs use caudoviricete ancestor-derived proteins (established either via significant similarity of at least some GTA proteins to caudeviricete proteins or by image-based evidence of caudovirion-like particles) to form caudovirion-like particles;
2. GTAs encapsidate mostly random pieces of host DNA (established experimentally);
3. GTA genomes are fully endogenized in host genomes, often across multiple loci (established experimentally and via genomic examination);
4. GTA genomes are not/cannot be fully packaged into particles due to limited particle head size (established via comparison of the packaged DNA length and size of GTA loci);
5. GTA genomes are mostly vertically inherited and GTAs co-diversify with their hosts (established via congruence between phylogenies of host and GTA genes); and
6. DNA encapsidated in GTA particles is delivered to other cells (established experimentally).

Having these attributes, GTAs have lost the ability to replicate and have become fully exapted by their cellular hosts. They are produced under specific conditions (e.g., nutrient depletion [Westbye et al. 2017b]) and mediate horizontal gene transfer (HGT), typically among cells of the same species.

The first GTA discovered, of the alphaproteobacterium *Rhodobacter capsulatus*, was described in 1974 by Barry Marrs (Marrs 1974). Since that time, distinct functional GTAs have been described in other alphaproteobacteria, a sulfate-reducing deltaproteobacterium, a methanogenic archaeon, and a spirochete that infects domestic pigs (Bertani 1999; Guy et al. 2013; Humphrey et al. 1997; Rapp and Wall 1987). Additionally, clusters of genes homologous to those encoding the *R. capsulatus* GTA are found in many alphaproteobacterial genomes, suggesting a wider prevalence of GTA production than presently appreciated (Kogay et al. 2019; Lang et al. 2002; Lang and Beatty 2007; Shakya et al. 2017). Indeed, some of these bacteria produce functional GTAs (Biers et al. 2008; Nagao et al. 2015; Tomasch et al. 2018).

The recent ICTV recognition of viriforms and the formal establishment of *Polydnaviriformidae* provides an opportunity to initiate a systematic classification of GTAs. Here we outline initial steps to establish such a formal taxonomic scheme for GTA viriforms, focusing specifically on GTAs experimentally documented as being produced by cells and performing gene transfer—and for which the genetic basis of particle production has been established. Simultaneously, we have also officially proposed this taxonomic scheme to the ICTV for the 2022–2023 proposal cycle.

## NOMENCLATURE OF GENE TRANSFER AGENTS AND ASSOCIATED TAXA

Per ICTV rules, virus names are written in lower case (except if a name component is a proper noun), without italics in any part of the name (even if a host species name is part of the name), and ending in the term “virus”, which in virus name abbreviations is “V”. Examples are measles virus (MeV) and Ebola virus (EBOV). The nomenclature of already classified viriforms (polydnaviriformids) follows these rules, with “virus” being replaced by “viriform” and the abbreviation “V” being replaced with “Vf” (e.g., “Glyptapanteles liparidis bracoviriform” is abbreviated “GlBVf”). We suggest applying these general rules to GTAs, but with “viriform” being replaced by “gene transfer agent” due to the long-established use of this phrase and “Vf” being replaced with “GTA”. Therefore, the gene transfer agent produced by *Rhodobacter capsulatus* would be called “Rhodobacter capsulatus gene transfer agent” and abbreviated as “RcGTA”, consistent with the established use of this abbreviation in the literature.

Rules for viriform taxon naming have been established by the ICVCN. Specifically,

> “[t]he formal endings for taxon names of viriforms are the suffixes “-*viriformia*” for realms, “-*viriforma*” for subrealms, “-*viriformae*” for kingdoms, “-*viriformites*” for subkingdoms, “-*viriformicota*” for phyla, “-*viriformicotina*” for subphyla, “-*viriformicetes*” for classes, “-*viriformicetidae*” for subclasses, “-*viriformales*” for orders, “-*viriformineae*” for suborders, “-*viriformidae*” for families, “-*viriforminae*” for subfamilies, and “-*viriform*” for genera and subgenera” (ICVCN Rule 3.26) (International Committee on Taxonomy of Viruses 2022; Kuhn et al. 2020)

and

> “[a] species name shall consist of only two distinct word components separated by a space. The first word component shall begin with a capital letter and be identical in spelling to the name of the genus to which the species belongs. The second word component shall not contain any suffixes specific for taxa of higher ranks. The entire species name (both word components) shall be italicized” (International Committee on Taxonomy of Viruses 2022).

We suggest adding the infix -*gta*-prior to the taxon-specific suffixes for immediate recognition of GTA-specific taxa (e.g., -*gtaviriform*).

## GENE TRANSFER AGENTS CAN BE ASSIGNED TO AT LEAST THREE MAJOR CLADES

Based on functionally and genetically characterized GTAs, at least three major GTA clades can be delineated.

### Alphaproteobacterial type I GTAs

The best characterized GTA of this clade is RcGTA, produced by *R. capsulatu*s (*Pseudomonadota*: *Alphaproteobacteria*: *Rhodobacterales*: *Rhodobacteraceae*). We designate RcGTA here as the founding member of one major GTA clade, the alphaproteobacterial type I GTAs. For many years since its discovery (Marrs 1974), RcGTA was the only known GTA. Now we know that homologous GTAs are produced by other bacteria from the order *Rhodobacterales*: *Dinoroseobacter shibae* (Dinoroseobacter shibae gene transfer agent [DsGTA]) (Tomasch et al. 2018), *Ruegeria pomeroyi* (Ruegeria pomeroyi gene transfer agent [RpGTA]) (Biers et al. 2008), and *Rhodovulum sulfidophilum* (Rhodovulum sulfidophilum gene transfer agent [RsGTA]) (Nagao et al. 2015). Additionally, genes encoding RcGTA-like GTAs are conserved in most genomes in the order *Rhodobacterales* and in many genomes of the alphaproteobacterial orders *Caulobacterales, Sphingomonadales, Parvibaculales*, and *Hyphomicrobiales* (formerly *Rhizobiales*) (Kogay et al. 2019; Lang et al. 2002; Lang and Beatty 2007; Shakya et al. 2017).

RcGTA and RcGTA-like GTA genes are similar in sequence to those of viruses classified in the uroviricot class *Caudoviricetes* (*Duplodnaviria*: *Heunggongvirae*) (Shakya et al. 2017). These GTAs are transmitted vertically from a bacterial parent to progeny during cell division (Lang and Beatty 2007; Shakya et al. 2017), similar to propagation of temperate viruses (“prophages”). However, in contrast to temperate virus genomes, the set of genes required for production of the GTA particle (the GTA “genome”) is not necessarily localized in one region of the host genome. In the case of RcGTA, known structural and regulatory genes are scattered across five loci in the *R. capsulatus* genome (Hynes et al. 2016), cumulatively spanning approximately 20 kilobases (kb) (**Figure 1** and **Supplementary Table S1**). Moreover, cellular regulatory genes are involved in controlling GTA particle production (Westbye et al. 2017a), adding another factor that makes the GTA genome difficult to differentiate from its host’s genome.

**Figure 1.**
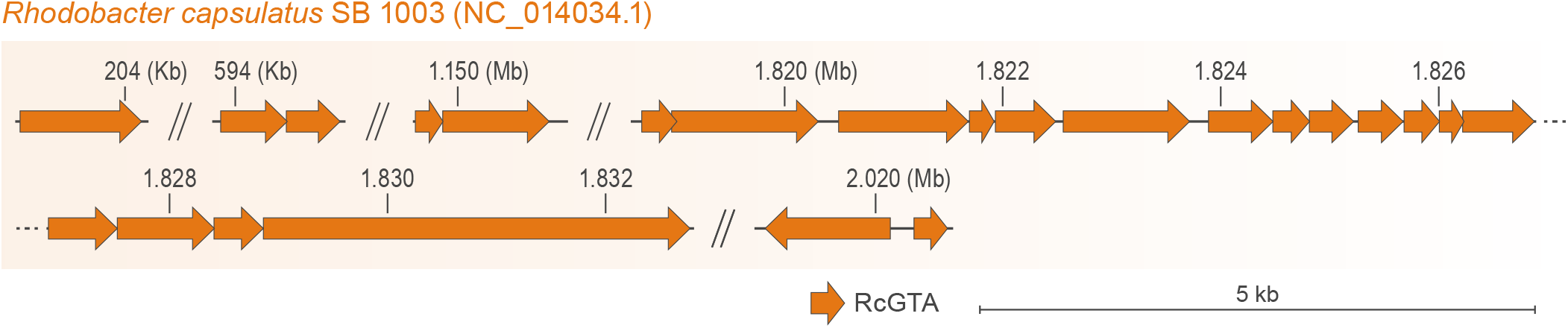
Genome of Rhodobacter capsulatus gene transfer agent (RcGTA). Genes (arrows) are depicted to scale, in their locations in the host genome (*R. capsulatus*). Exact coordinates of the RcGTA genes, their locus tags, and their functional annotations are listed in **Supplementary Table S1**.

RcGTA particles resemble virions of caudoviricetes (Yen et al. 1979) and have been structurally characterized at high resolution (Bárdy et al. 2020). RcGTA particles have head diameters of 38 nm and tail lengths of 49 nm. A small percentage of RcGTA particles have T = 3 quasi-icosahedral heads, but the capsid shape of most particles is oblate, as they lack the five hexamers of capsid protein needed to form genuine icosahedral heads. Because of the small head size, RcGTA particles can only package double-stranded DNA of approximately 4 kb in length (Yen et al. 1979). The DNA is also encapsidated at 10–25% lower density than typical caudoviricetes (Bárdy et al. 2020). Both RcGTA particle production and acquisition of the GTA-packaged DNA by other host cells in the population are controlled by the same cellular regulatory systems (Westbye et al. 2017a). Only 0.1–3.0% of cells produce GTA particles (Fogg et al. 2012; Hynes et al. 2012), whereas the remaining cells produce a GTA receptor (Brimacombe et al. 2013).

Compositionally, structural proteins encoded by RcGTA and RcGTA-like GTAs are biased towards amino acids that are energetically cheaper to produce (Kogay et al. 2020). To date, such a bias has not yet been associated with viruses. Based on this difference in amino-acid composition, GTA proteins can be distinguished from their viral homologs using a machine-learning approach, which is implemented in the publicly available GTA-Hunter program (Kogay et al. 2019).

In a comprehensive evolutionary analysis of homologs of the large subunit of the DNA packaging terminase enzyme (TerL, encoded by the *g2* gene in the RcGTA genome), RcGTA and RcGTA-like GTAs form a clade closely related to, but distinct from, duplodnavirians (Esterman et al. 2021). To illustrate the relationships of alphaproteobacterial type I GTAs to each other and to their closest viral homologs, we reconstructed evolutionary histories of their TerL proteins and the HK97-like major capsid proteins (HK97-MCP, encoded by the *g5* gene in the GTA genome, is the hallmark protein that defines the virus realm *Duplodnaviria* [Koonin et al. 2020]). Consistent with an earlier analysis (Esterman et al. 2021), RcGTA and RcGTA-like GTAs formed a clade closely related to, but distinct from, caudoviricetes (**Figure 2**), with a few exceptions that are likely artefacts of phylogenetic reconstruction.

**Figure 2.**
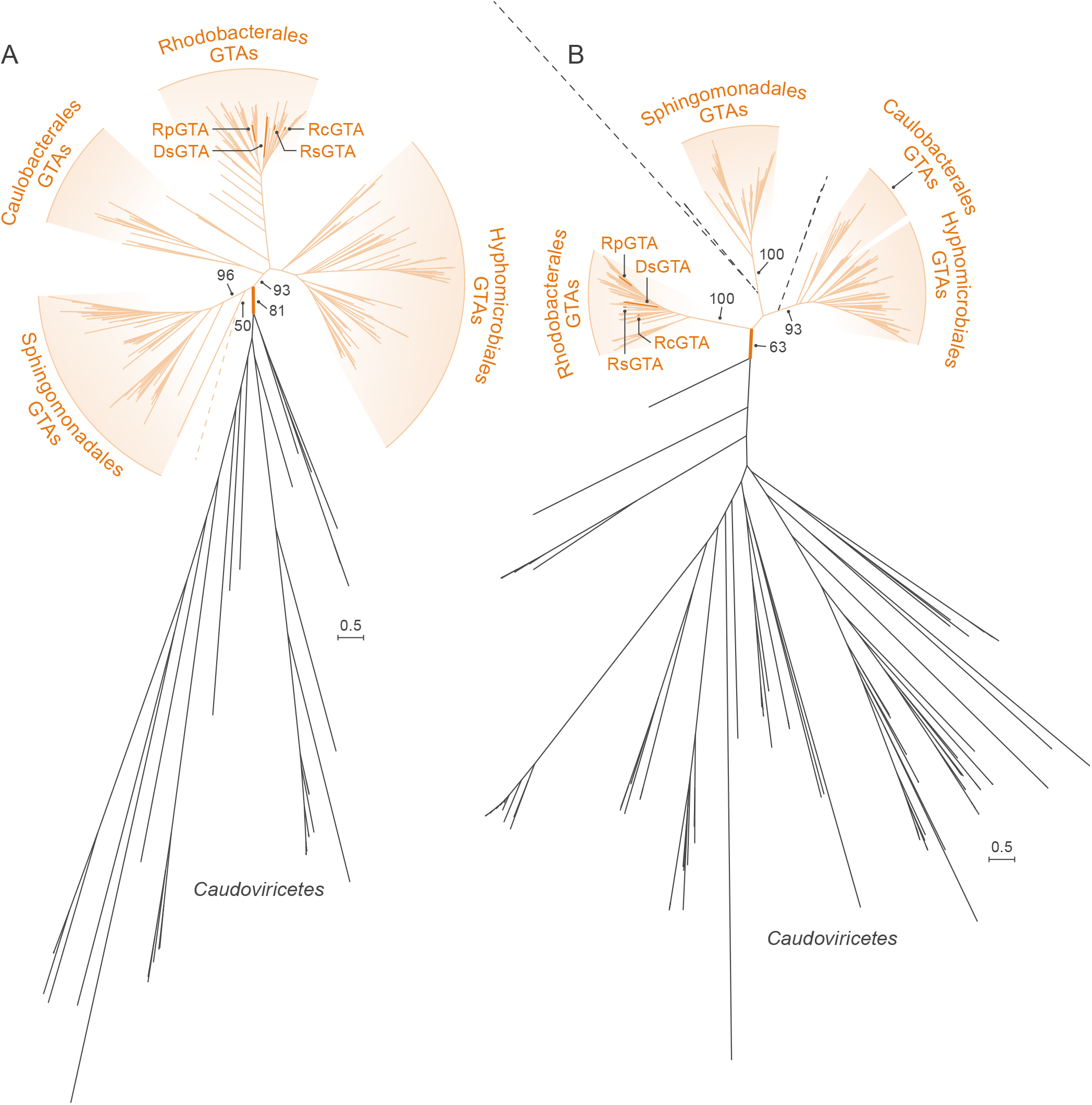
Maximum Likelihood phylogenies of (A) large terminase (TerL) subunits and (B) HK97 major capsid protein (HK97-MCP) sequences of rhodogtaviriformids and their closest known caudoviricete homologs. Alphaproteobacterial type I gene transfer agent (GTA) (rhodogtaviriformid) lineages are shown in orange. Caudoviricete lineages that are nested within GTA lineages are shown in dashed black lines. Other caudoviricete lineages are shown in solid black lines. Bootstrap support values are shown only for selected branches. Scale bars represent substitutions per site. DsGTA, Dinoroseobacter shibae gene transfer agent; GTA, gene transfer agent; RcGTA, Rhodobacter capsulatus gene transfer agent; RpGTA, Ruegeria pomeroyi gene transfer agent; RsGTA, Rhodovolum sulfidophilum, gene transfer agent.

Specifically, in the TerL phylogeny (**Figure 2A**), all viral homologs except one (Caulobacter virus Sansa) are separated from GTA proteins (with a solid bootstrap support of 81%). Caulobacter virus Sansa groups with one GTA sequence from a bacterium of the order *Sphingomonadales* (with a low bootstrap support of 50%), whereas all other GTAs of *Sphingomonadales* bacteria group together (with a strong bootstrap support of 96%). We hypothesize that the phylogenetic placement of the Caulobacter virus Sansa TerL is due to the long-branch attraction artefact (Felsenstein 1978). We searched for a maximum-likelihood tree in which caudoviricete- and GTA-derived TerLs were required to group separately from each other and compared that tree to the tree depicted in **Figure 2A**. We found that the likelihoods of the two trees are not significantly different (approximately unbiased [AU] test; p-value = 0.555), confirming that the placement of the Caulobacter virus Sansa sequence within the GTA sequences is unreliable.

In the HK97-MCP phylogeny (**Figure 2B**), GTAs and most caudoviricetes are separated by a branch with 63% bootstrap support. Several caudoviricetes that group within GTAs are located on long branches, are situated outside of well-supported groups of GTAs from several alphaproteobacterial orders and have very low bootstrap support for their placements. It is therefore likely that the positions of these viral homologs are unreliable. To test this hypothesis, we identified a maximum-likelihood phylogeny among trees in which GTAs and caudoviricetes were required to be separated by a branch. The likelihoods of this tree and the phylogeny shown in **Figure 2B** are not significantly different (AU test; p-value = 0.534). Therefore, these viruses are likely positioned in different places in trees reconstructed from different bootstrap replicates, which would lead to their artificial (and poorly supported) basal positions with the GTA homologs on the tree shown in **Figure 2B**.

In the **Figure 2** trees, GTA branches have shorter lengths than their caudoviricete counterparts, conforming with the reported slower evolutionary rate of GTAs compared to viruses (Shakya et al. 2017). Additionally, on both phylogenetic trees, GTAs from alphaproteobacteria of different orders form separate groups with very high support, corroborating vertical inheritance of most GTA genes (Lang et al. 2002; Lang and Beatty 2007; Shakya et al. 2017).

Together, these results justify the classification of RcGTA and three RcGTA-like GTAs in a common viriform taxon: family *Rhodogtaviriformidae* (from *Rhodobacterales*, infix -*gta*-, and family-specific suffix -*<u>viriformidae</u>*). Given limited dataset size (i.e., just four GTAs), it is challenging to establish quantifiable criteria for demarcating taxonomic relationships among the four GTAs. In the future, when more GTA sequences become available for analyses, a criterion based on percent sequence similarity among shared genes should be considered. For now, based on the evidence of co-evolution of these GTAs and their specific hosts, we argue that at least four rhodogtaviriformid genera, each for GTAs of bacteria classified in distinct genera included in *Rhodobacterales*, ought to be established:

- *Dinogtaviriform* (named after DsGTA host genus *Dinoroseobacter*, infix -*gta*-, and genus-specific suffix -*<u>viriform</u>*) to include one new species, *Dinogtaviriform tomaschi* (species epithet to honor GTA researcher Jürgen <u>Tomasch</u>, who was instrumental in the discovery of DsGTA) for DsGTA (**Supplementary Table S2**);
- *Rhodobactegtaviriform* (named after RcGTA host genus <u>Rhodobacter</u>, infix -*<u>gta</u>*-, and genus-specific suffix -*<u>viriform</u>*) to include one new species, *Rhodobactegtaviriform marrsi* (species epithet to honor GTA researcher Barry <u>Marrs</u>, who first discovered GTAs and coined the term “gene transfer agent”) for RcGTA (**Supplementary Table S1**);
- *Rhodovulugtaviriform* (named after RsGTA host genus *Rhodovulum*, infix -*gta*-, and genus-specific suffix -*<u>viriform</u>*) to include one new species, *Rhodovulugtaviriform kikuchii* (species epithet to honor GTA researcher Yo <u>Kikuchi</u>, who was instrumental in the discovery of RsGTA) for RsGTA (**Supplementary Table S3**); and
- *Ruegerigtaviriform* (named after RpGTA host genus *Ruegeria*, infix -*<u>gta</u>*-, and genus-specific suffix -*viriform*) to include one new species, *Ruegerigtaviriform cheni* (species epithet to honor GTA researcher Feng <u>Chen</u>, who was instrumental in the discovery of RpGTA) for RpGTA (**Supplementary Table S4**).

### Alphaproteobacterial type II GTAs

There was a lag between discovery of these elements and their recognitions as bona fide GTAs. Phage-like particles, originally referred to as bacteriophage-like particles (BLPs), that contained heterogenous DNA from *Bartonella* host genomes were first characterized in *B. henselae* (Anderson et al. 1994), and noted to be similar in structure to particles produced by *B. bacilliformis* (Umemori et al. 1992). These *B. bacilliformis* particles were subsequently shown to also contain heterogeneous genomic DNA fragments, but attempts to demonstrate their gene transfer ability were not successful (Barbian and Minnick 2000). Functionality of the particles produced by *Bartonella* for gene transfer (Bartonella gene transfer agent [BaGTA]) was eventually demonstrated by work on *B. henselae* (*Pseudomonadota*: *Alphaproteobacteria*: *Hyphomicrobiales*: *Bartonellaceae*) (Guy et al. 2013). BaGTA genes were initially proposed to be located within a single cluster of 11–13 genes spanning approximately 14 kb (Guy et al. 2013). However, a subsequent screen for genes essential for BaGTA functionality identified a total of 29 genes located within a larger (approximately 79-kb-long) region (Québatte et al. 2017) (**Figure 3** and **Supplementary Table S5**). Homologs of BaGTA genes (BaGTA-like GTAs) were found in the genomes of multiple species of *Bartonella* (Berglund et al. 2009; Guy et al. 2013; Tamarit et al. 2018). BaGTA genes are located near an active virus-derived origin of replication and next to genes encoding secretion systems (Guy et al. 2013). As a result, the region of the genome containing BaGTA and these secretion-system genes are amplified and packaged more often than other genomic regions (Guy et al. 2013; Québatte et al. 2017). These findings led to the hypothesis that BaGTA and BaGTA-like GTAs have been maintained due to their mediation of HGT of secretion-system and toxin genes, thereby enabling *Bartonella* bacteria to adapt to diverse hosts (Guy et al. 2013). However, actual GTA-mediated DNA transfer among bacterial cells has only been demonstrated for *B. henselae* (Guy et al. 2013).

**Figure 3.**
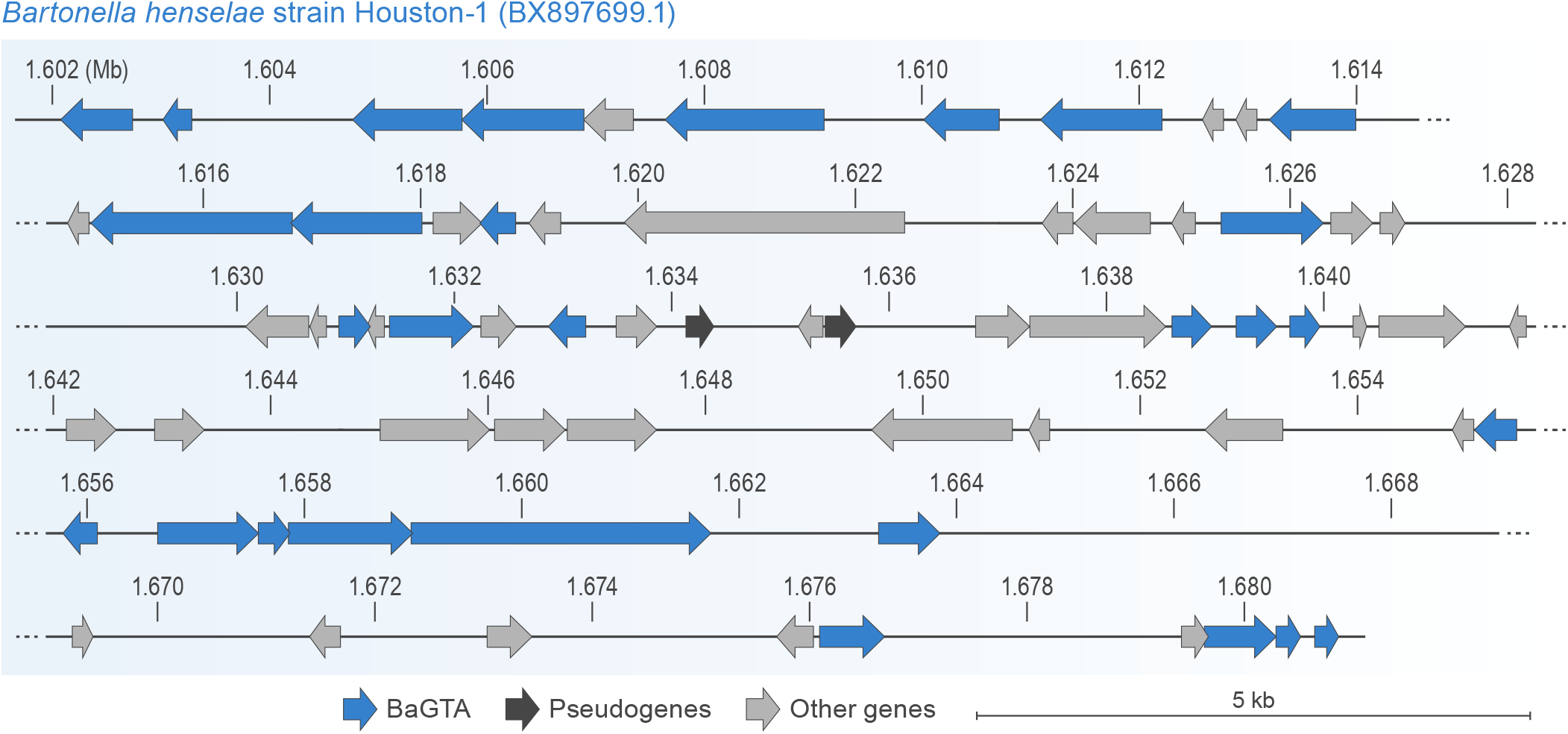
Genome of Bartonella gene transfer agent (BaGTA). Genes (arrows) are depicted to scale, in their locations in the host genome (*B. henselae*). Exact coordinates of the BaGTA genes, their locus tags, and their functional annotations are listed in **Supplementary Table S5**.

There, BaGTA production is restricted to a distinct subpopulation of fast-growing cells, which comprise about 6% of the total population (Québatte et al. 2017), and the uptake of BaGTA-packaged DNA was proposed to be limited to cells undergoing division (Québatte et al. 2017).

There are some discrepancies in the literature regarding the structure of BaGTA particles, suggesting some bacteria might release additional phage-like particles. The *B. henselae* particles were originally reported as particles without tails or with short non-contractile tails with a head diameter of 40 nm (Anderson et al. 1994). The head diameter of the *B. bacilliformis* particles was originally measured at 40 nm (Umemori et al. 1992) and subsequently 80 nm (Barbian and Minnick 2000). Those of *B. grahamii* were reported as possessing long non-contractile tails and icosahedral heads of 50–70 nm or 80 nm and tails of 100 nm (Berglund et al. 2009). Although BaGTA particles are potentially able to package the entire main structural gene cluster of 11–13 genes, they cannot package all 29 genes required for BaGTA production due to a capacity of 14 kb (Anderson et al. 1994; Guy et al. 2013; Lang et al. 2017).

In the TerL phylogeny, BaGTA-like homologs are separated from almost all caudoviricetes by longer branches (with 100% bootstrap support; **Figure 4A**). Two caudoviricete homologs (Sulfitobacter phage pCB2047-C and Sulfitobacter phage NYA-2014a) group together and are nested within the BaGTA-like group (with 84% bootstrap support). We hypothesize that the *terL* gene was horizontally transferred from GTAs to these caudoviricetes, with similar HGT events documented between RcGTA-like GTAs and caudoviricetes infecting bacteria of the *Rhodobacterales* (Zhan et al. 2016). In the HK97-MCP phylogeny, BaGTA homologs are located on shorter branches than their caudoviricete counterparts and are separated from caudoviricete homologs with 100% bootstrap support (**Figure 4B**). Phylogenomic analyses suggest that *Bartonella* GTAs have co-evolved with their hosts (Tamarit et al. 2018).

**Figure 4.**
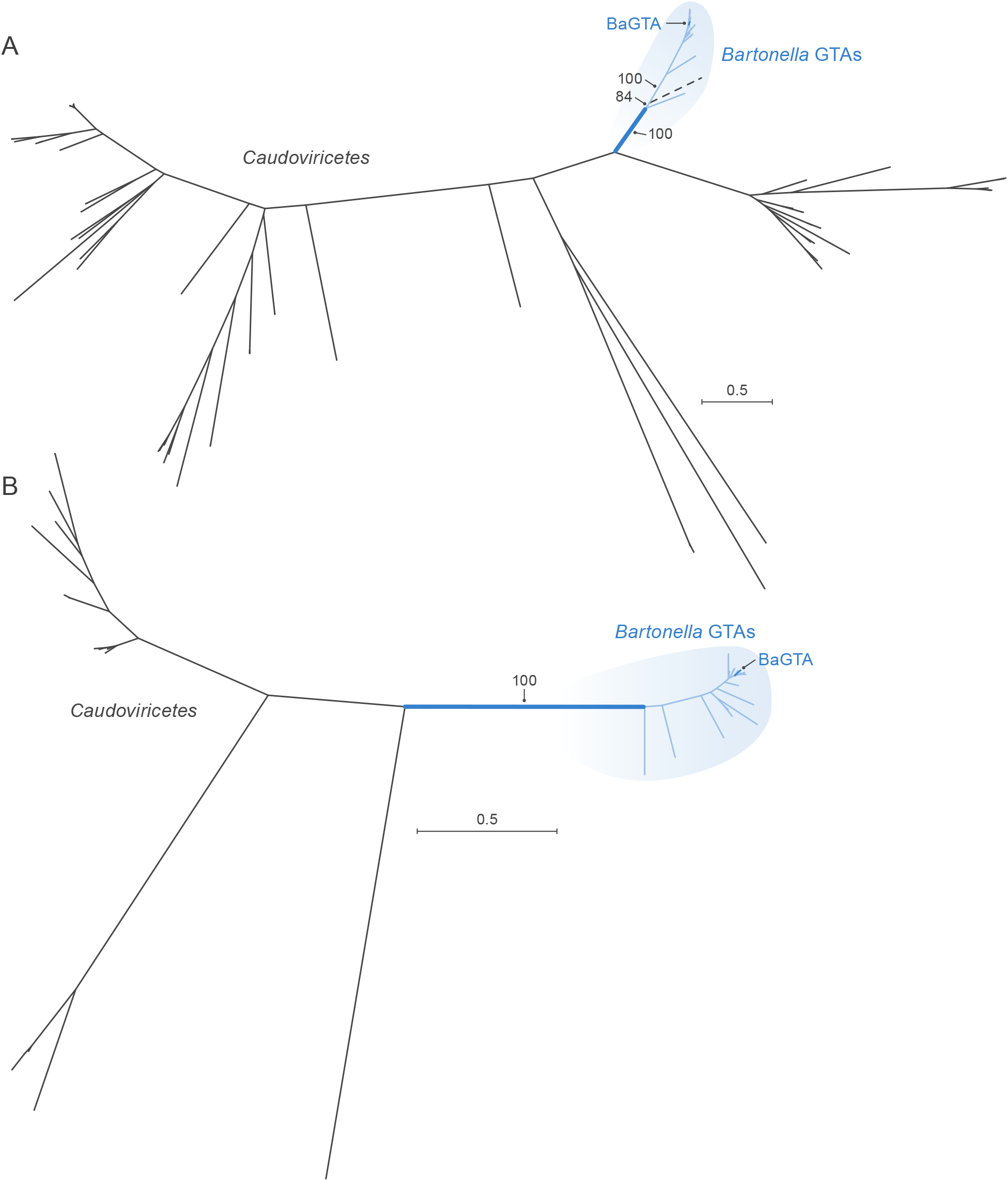
Maximum Likelihood phylogenies of (A) large terminase (TerL) subunits and (B) HK97 major capsid protein (HK97-MCP) sequences of bartogtaviriformids and their closest known caudoviricete homologs. Alphaproteobacterial type II gene transfer agent (GTA) (bartogtaviriformid) lineages are shown in blue. Caudoviricete lineages are shown in black. Two nearly identical caudoviricete lineages that are nested within GTA lineages are shown in dashed black lines. A bootstrap support value is shown only for the branch separating GTA and caudoviricete sequences. Scale bars indicate substitutions per site. BaGTA, Bartonella gene transfer agent; GTA, gene transfer agent.

Together, these results justify the classification of BaGTA and BaGTA-like GTAs in a common viriform taxon, family *Bartogtaviriformidae* (from *Bartonella*, infix -*<u>gta</u>*-, and family-specific suffix -*<u>viriformidae</u>*). For now, we argue that at least one bartogtaviriformid genus ought to be established: *Bartonegtaviriform* (named after BaGTA host genus *Bartonella*, infix -*gta*-, and genus-specific suffix -*<u>viriform</u>*) including one new species, *Bartonegtaviriform andersoni* (species epithet to honor GTA researcher Burt <u>Anderson</u>, who first discovered BaGTA particles [Anderson et al. 1994]) for BaGTA.

### GTAs of spirochaetes

A GTA originally called virus of *Serpulina hyodysenteriae* 1 (VSH-1) was identified in *Brachyspira* (formerly *Serpulina*) *hyodysenteriae* (*Spirochaetota*: *Spirochaetia*: *Brachyspirales*: *Brachyspiraceae*) (Humphrey et al. 1997). In accordance with the nomenclature rules established here, we suggest renaming this GTA to Brachyspira hyodysenteriae gene transfer agent (BhGTA). The structural gene cluster responsible for production of BhGTA particles—i.e., the BhGTA “genome”—is 16.3 kb in length (Matson et al. 2005) (**Figure 5** and **Supplementary Table S6**).

**Figure 5.**
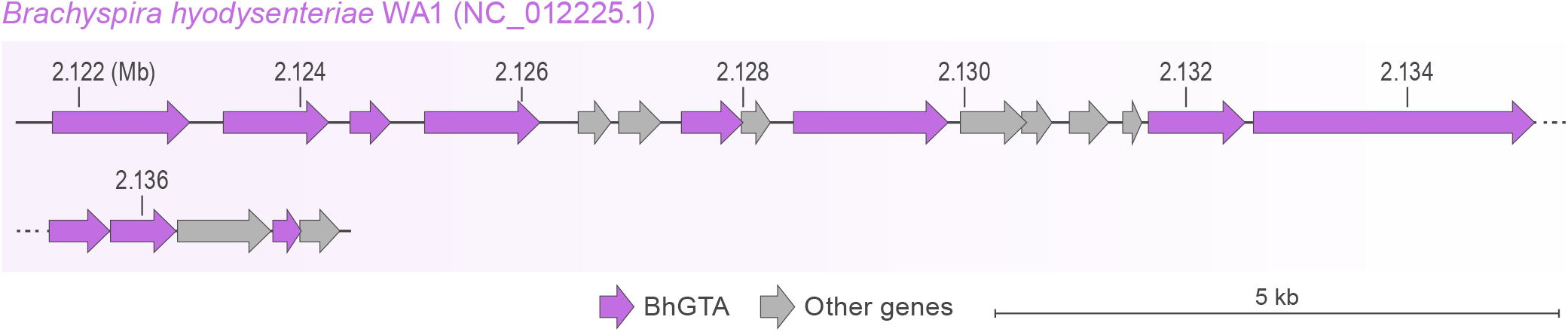
Genome of Brachyspira hyodysenteriae gene transfer agent (BhGTA). Genes (arrows) are depicted to scale, in their locations in the host genome (*B. hyodysenteriae*). Exact coordinates of the BhGTA genes, their locus tags, and their functional annotations are listed in **Supplementary Table S6**.

BhGTA particles have a head diameter of 45 nm and a flexible non-contractile tail of 65 nm (Humphrey et al. 1997). Like other GTAs, BhGTA is unable to package and transfer its entire genome, given the limiting capacity of 7.5 kb (Humphrey et al. 1997; Matson et al. 2005). Restriction enzyme digests of the packaged DNA and the range of marker genes that can be transferred by BhGTA particles suggest that they package any region of the *B. hyodysenteriae* genome (Humphrey et al. 1997) without an obvious bias for the genomic region that encodes BhGTA. The induction of BhGTA particle production by DNA-damaging agents, such as mitomycin C and antibiotics, results in large-scale lysis of cells (Stanton et al. 2008). However, the proportion of *B. hyodysenteriae* cells in a population that naturally produce and release BhGTA particles has not been quantified. BhGTA particles are capable of transferring antimicrobial resistance genes within the bacterial population (Stanton et al. 2008), pointing at possible selective advantages of maintaining the capability of BhGTA particle production.

Homologs of genes in the BhGTA genome were found in the genomes of other members of the genus *Brachyspira*, but there is no gene synteny in their organization (Motro et al. 2009). Unlike in rhodogtaviriformids and bartogtaviriformids, an endolysin-encoding gene is the only gene in the BhGTA genome that has a significant sequence similarity to caudoviricete genes in the National Center for Biotechnology Information (NCBI) Reference Sequence (RefSeq) database (accessed in May 2022). Some genes encoding the BhGTA particle proteins were experimentally validated (including endolysin), and the particles structurally resemble those of caudoviricetes (Matson et al. 2005). Therefore, the absence of their homologs in the viral RefSeq database is likely due to the limited sampling of the virosphere.

In the endolysin phylogeny, the *Brachyspira* homologs group together and are separated from all caudoviricetes by a long branch (with 100% bootstrap support; **Figure 6A**). Additionally, the *B. hyodysenteriae* genome encodes a single copy of an identifiable *terL* gene, which is located outside of the currently delineated BhGTA genome. Homologs of this *terL* gene are also present in a single copy in genomes of other *Brachyspira* bacteria that encode BhGTA-like MCPs. These homologs are highly conserved, with pairwise amino-acid identities of 81– 100%. In a phylogenetic tree, the *Brachyspira* TerLs are separated from all caudoviricete TerLs by a longer branch (with 100% bootstrap support; **Figure 6B**). Although the role of this TerL homolog in the BhGTA lifecycle has not been experimentally validated, the presence of the encoding gene as the only identifiable *terL* in the *Brachyspira* genomes, its high degree of conservation within the *Brachyspira* genus and its divergence from the related caudoviricete sequences support its potential involvement in the packaging of DNA into the BhGTA particles. Based on comparison of *Brachyspira* GTA and host genes, GTAs have co-diversified with *Brachyspira* (Motro et al. 2009).

**Figure 6.**
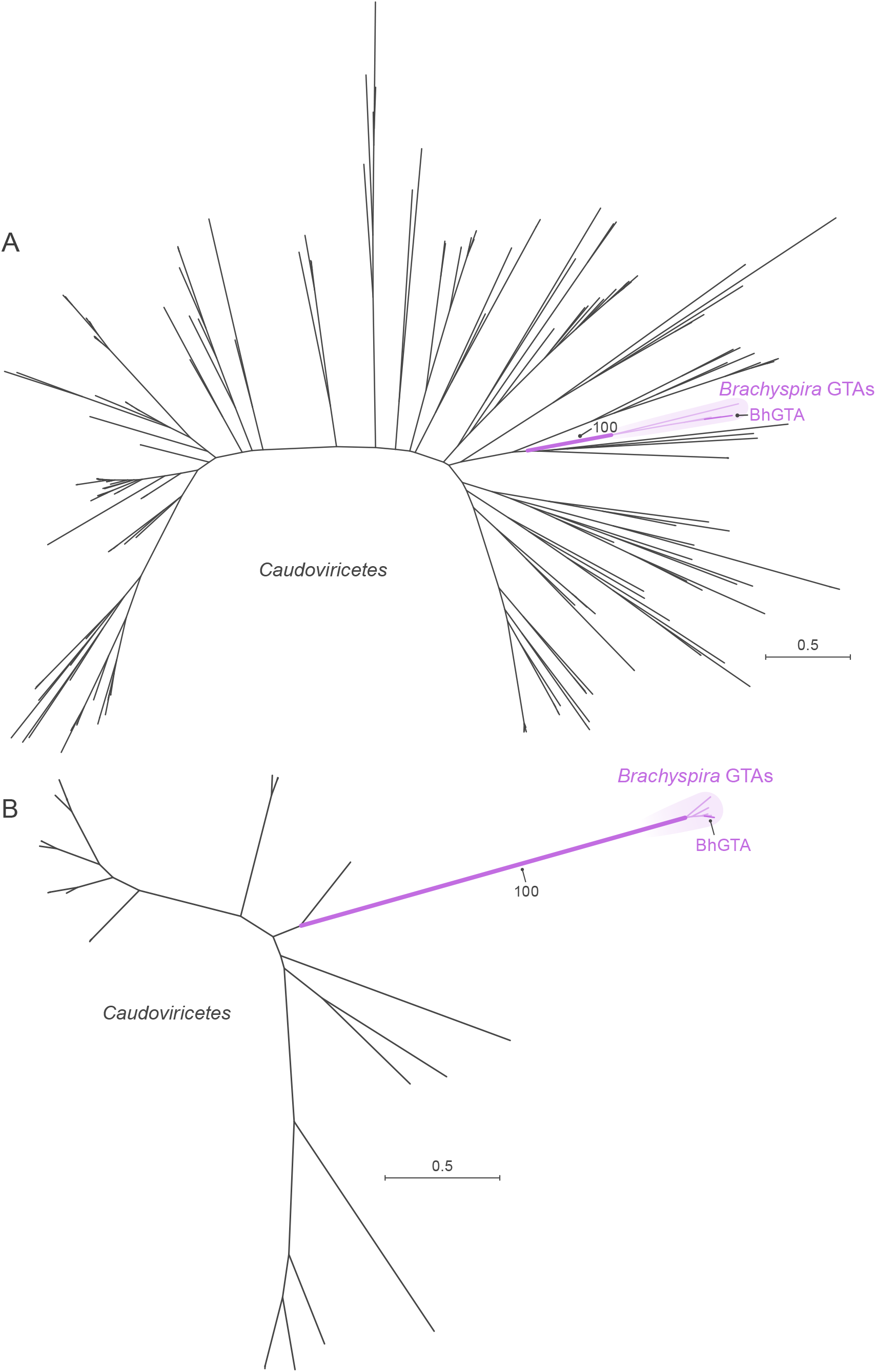
Maximum Likelihood phylogenies of (A) endolysin and (B) the putative large terminase (TerL) subunits of brachygtaviriformids and their closest known caudoviricete homologs. *Brachyspira* gene transfer agent (GTA) lineages are shown in purple. Caudoviricete lineages are shown in black. A bootstrap support value is shown only for the branch separating GTA and caudoviricete sequences. Scale bar indicates substitutions per site. BhGTA, Brachyspira hyodysenteriae gene transfer agent; GTA, gene transfer agent.

Together, these results justify the classification of BhGTA and BhGTA-like GTAs in a common viriform taxon, family *Brachygtaviriformidae* (form *Brachyspira*, infix -*<u>gta</u>*-, and family-specific suffix -*<u>viriformidae</u>*). For now, we argue that at least one brachygtaviriformid genus ought to be established: *Brachyspigtaviriform* (named after BhGTA host genus *Brachyspira*, infix -*gta*-, and genus-specific suffix -*<u>viriform</u>*) to include one new species, *Brachyspigtaviriform stantoni* (species epithet to honor GTA researcher Thaddeus Stanton, who first discovered BhGTA particles [Humphrey et al. 1997]) for BhGTA.

### Independent origins of the three GTAs

Genes from the genomes of these three GTAs are either not homologous or too divergent to have significant sequence similarity in BLASTP searches of the encoded proteins. For example, pairwise amino-acid identity of TerLs, which is one of the most conserved GTA and caudoviricete proteins, is 14–20% among RcGTA, BaGTA, and BhGTA. Nevertheless, an iterative clustering-alignment-phylogeny procedure (Wolf et al. 2018) established the homology among known TerL proteins that include RcGTA, BaGTA, and putative BhGTA TerLs (Esterman et al. 2021). The evolutionary history of RcGTA-like, BaGTA-like, putative BhGTA-like TerLs, and their closest known caudoviricete homologs (**Figure 7**) demonstrates that GTA-like TerLs appear in three distinct clades within viral TerLs. Based on this phylogenetic evidence, we propose that these three GTA clades are a result of three independent exaptation events. Therefore, just like viruses (which are classified in at least six unrelated realms), GTA viriforms are polyphyletic.

**Figure 7.**
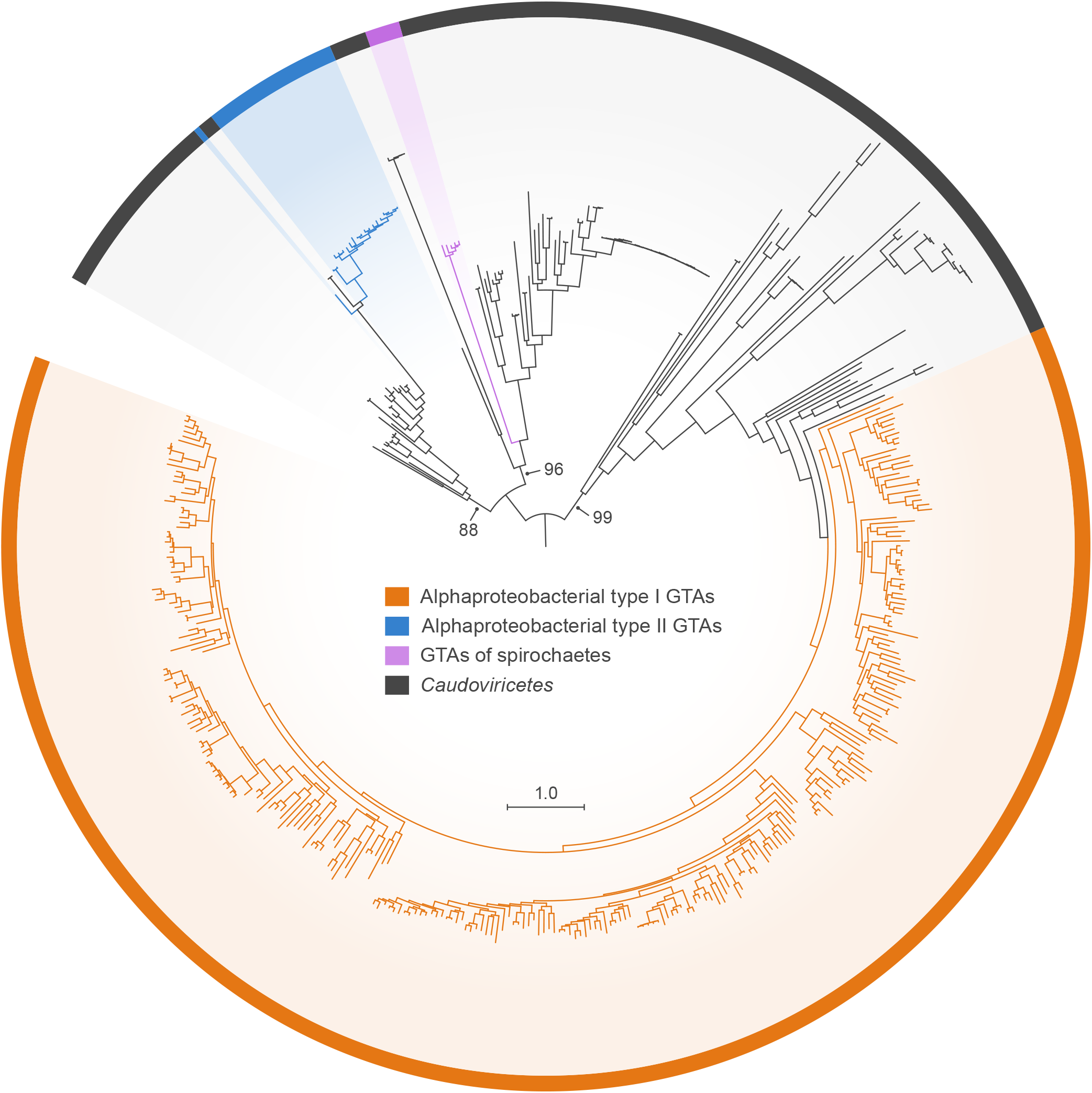
Maximum Likelihood phylogeny of the large terminase (TerL) subunits of three major clades of GTAs and their closest known caudoviricete homologs. This tree includes all TerL homologs from Figures 2A, 4A, and 6A, using their color coding. Bootstrap support values are shown only for the branches separating three GTA clades and their closest caudoviricete sequences. Scale bar indicates substitutions per site. GTA, gene transfer agent.

## DISCUSSION

Based on the evolutionary differences between GTA and caudoviricete genes encoding well-conserved proteins and on morphological differences of GTA particles, we propose three families for these GTAs. The greatest number of functionally confirmed and putative GTAs are in the alphaproteobacterial type I clade, which, for now, is proposed to be a family *Rhodogtaviriformidae* that includes at least four genera. The members of this family are currently restricted to a single cellular order (*Rhodobacterales*). The TerLs and MCPs of these RcGTA-like GTAs and alphaproteobacterial type II GTAs (*Bartogtaviriformidae*) are clearly distinguishable from each other and their caudoviricete homologs and evolve at a slower rate (**Figure 2** and **Figure 4**) (Esterman et al. 2021; Shakya et al. 2017). The spirochaete GTAs (*Brachygtaviriformidae*) are more difficult to distinguish from caudoviricetes due to a lack of available viral representatives in GenBank for all but one experimentally validated BhGTA gene. Nevertheless, both the experimentally validated BhGTA endolysin and the putative BhGTA TerL and their *Brachyspira* homologs also form a well-supported cluster distinct from caudoviricete lineages; moreover, brachygtaviriformid TerLs evolve at a slower rate than their spirochete homologs (**Figure 6B**). As in the case with the experimentally validated RcGTA, the “genome” of BhGTA is also likely dispersed across multiple loci.

Analyses of environmental samples and genome sequences suggest the existence of a large number of GTAs, especially those related to the rhodogtaviriformids (Biers et al. 2008; Fu et al. 2010; McDaniel et al. 2010; Zhao et al. 2009). In a genome-wide screen of 1,423 alphaproteobacterial genomes, 57.5% were found to encode RcGTA-like “genomes”, which are often annotated as either intact or incomplete prophages (Kogay et al. 2019). The great majority of RcGTA-like genes in alphaproteobacterial genomes are associated with bacteria for which a GTA-based gene-transfer activity has not been documented, and it is possible that some of these RcGTA-like genes may not be expressed to produce functional particles. Therefore, we have restricted our proposal to those GTAs that have been shown to be functional. However, we speculate that at least some (and perhaps many) of these GTA-like gene clusters will be shown to produce functional GTAs that will need to be classified.

Based on the evolutionary history of TerL proteins (**Figure 7**), it is likely that the proposed three GTA families had distinct caudoviricete progenitors. Eventual deduction of the relatives of these progenitors may make it possible (or necessary) to include these GTA families in the virus class *Caudoviricetes*, thereby creating an overarching taxon for distinct MGEs (viruses and viriforms). Since the exaptation events, however, the three families have evolved as part of the host genomes (Esterman et al. 2021; Lang et al. 2002; Lang and Beatty 2007; Shakya et al. 2017), in the case of the rhodogtaviriformids for hundreds of millions of years (Shakya et al. 2017). As a result, GTAs effectively became a component of cellular genomes, integrated into cellular regulatory circuits that also control processes such as motility, quorum sensing, extracellular polysaccharide synthesis, and biofilm formation (Lang et al. 2017; Pallegar et al. 2020; Shimizu et al. 2022). There is also mounting evidence that GTA genes experience selective pressures to be maintained in their host genomes (Kogay et al. 2020; Lang et al. 2012). Although the fitness benefits associated with GTA production remain to be elucidated, the time is now ripe to have the known GTAs officially recognized and classified as specific viriforms. We recognize this step as the initiation of a taxonomic framework that undoubtedly will rapidly expand and change in the future.

## METHODS

To identify alphaproteobacterial type I GTAs, we searched for RcGTA-like sequences in 1,248 complete alphaproteobacterial genomes extracted from the NCBI RefSeq database (accessed in October 2020) using GTA-Hunter (Kogay et al. 2019). We identified 503 genomes that contained at least six RcGTA homologs in the same genetic neighborhood and had both *g2* (encoding TerL) and *g5* (encoding HK97-MCP) genes. To remove redundancy, we clustered genomes into the operational taxonomic units (OTUs) using an average nucleotide identity threshold of 95%. From all genomes within an OTU, we selected one genome with the largest number of GTA genes. This strategy resulted in 290 representative GTAs selected for further analysis. We identified the closest viral homologs of the TerL and HK97-MCP proteins from these GTAs by conducting a BLASTP search (Altschul et al. 1997) of the RefSeq database (accessed in March 2021) (O’Leary et al. 2016), using TerL and HK97-MCP proteins from representative GTAs as queries, an e-value cutoff of 0.001, and query coverage of at least 50%. Retrieved viral homologs with identical amino-acid sequences were removed from further analyses. For both proteins, we aligned amino-acid sequences of GTA and virus homologs using MAFFT v7.455 with -linsi option (Katoh and Standley 2013). We reconstructed phylogenetic trees using IQ-TREE v2 (Minh et al. 2020), identifying the best substitution models using the built-in ModelFinder (Kalyaanamoorthy et al. 2017). The selected models were LG+F+R9 and LG+F+R7 for TerL and HK97-MCP datasets, respectively. Branch support values were assessed using 1,000 ultrafast bootstrap replicates and a hill-climbing nearest-neighbor interchange search for optimal trees (Hoang et al. 2018). Additionally, for both protein phylogenies, we reconstructed a phylogenetic tree in IQ-TREE v2 (Minh et al. 2020) using a tree search that was constrained by requiring all GTAs and all viruses to be separated by a branch. We compared the resultant trees in unconstrained and constrained searches using the AU test (Shimodaira 2002), as implemented in the IQ-TREE v2 program.

To identify alphaproteobacterial type II GTAs, we used the BaGTA TerL and HK97-MCP sequences (accession numbers WP_034448260.1 and WP_011181178.1, respectively) as queries in a BLASTP search against the 57 complete *Bartonella* genomes extracted from the RefSeq database (accessed in May 2022). We restricted our search only to matches for which BaGTA TerL and HK97-MCP homologs are in the same genomic neighborhood (defined as being within 5 kb of each other). In genomes with multiple matches to the query protein, we retained only the homolog with the highest BLASTP bit score. We clustered 57 genomes using a 95% average nucleotide identity (ANI) threshold and randomly selected one TerL and HK97-MCP representative from each cluster for phylogenetic analysis. We identified caudoviricete homologs by conducting a BLASTP search (e-value cutoff of 0.001, and query coverage of at least 50%) against viral RefSeq database (accessed in May 2022). We performed phylogenetic reconstructions as described above for alphaproteobacterial type I GTAs. The selected best substitution models were LG+R6 and LG+G4 for TerL and HK97-MCP datasets, respectively.

To identify GTAs of spirochaetes, we used BhGTA’s MCP sequence (GenBank accession number WP_012671344.1) as a query in a BLASTP search (with an e-value cutoff of 0.001 and query coverage of at least 50%) against the 13 complete *Brachyspira* genomes extracted from the RefSeq database (accessed in May 2022). We used TerL of *B. hyodysenteriae* (GenBank accession number WP_012671469.1) and endolysin protein of *B. hyodysenteriae* (GenBank accession number WP_012671356.1) as queries in a BLASTP search (with an e-value cutoff of 0.001 and query coverage of at least 50%) against the same set of 13 genomes. For endolysins, we only retained matches that co-localized within the BhGTA region on the chromosome. We clustered 13 genomes using a 95% ANI threshold and randomly chose one TerL and endolysin representative from each cluster for phylogenetic analyses. We identified caudoviricete homologs by doing BLASTP searches (e-value cutoff of 0.001 and query coverage of at least 50%) against the viral RefSeq database (accessed in May 2022). We performed phylogenetic reconstructions as described above for the alphaproteobacterial type I GTAs. The selected best substitution models were VT+F+R3 and WAG+R6 for TerL and endolysin datasets, respectively.

To reconstruct the phylogeny that includes all three clades of GTAs, we combined all TerL homologs extracted in the above-described procedures into one dataset. We aligned the TerL sequences using MAFFT v7.455 with -dash option (Rozewicki et al. 2019) and trimmed the obtained alignment using ClipKIT with -gappy option (Steenwyk et al. 2020). We computed the phylogenetic tree using IQ-TREE v2 (Minh et al. 2020) as described above with the LG+F+R10 substitution model selected by ModelFinder. We rooted the tree using a larger TerL phylogeny presented in Esterman et al. (2021).

We visualized all phylogenetic trees in iTOL v6 (Letunic and Bork 2021).

## ACKNOWLEDGEMENTS

We thank Anya Crane and Jiro Wada (Integrated Research Facility at Fort Detrick, Division of Clinical Research, National Institute of Allergy and Infectious Diseases, National Institutes of Health) for editing the manuscript and figures, respectively.

## SUPPLEMENTARY DATA

Amino-acid sequence alignments in FASTA format and phylogenetic trees in Newick format for trees in **Figures 2, 4, 6**, and **7** are available in **Supplementary_Data.zip** file.

## DATA AVAILABILITY

All data used in this manuscript were retrieved from publicly available GenBank databases, as described in the Methods. The accession numbers of database records used in the phylogenetic analyses can be found in alignments that are included in the **Supplementary Data**.

## FUNDING

This work was supported in part through Laulima Government Solutions, LLC, prime contract with the U.S. National Institute of Allergy and Infectious Diseases (NIAID) under Contract No. HHSN272201800013C; J.H.K. performed this work as an employee of Tunnell Government Services (TGS), a subcontractor of Laulima Government Solutions, LLC, under Contract No. HHSN272201800013C. This work was also supported by the Simons Foundation Investigator in Mathematical Modeling of Living Systems award #327936 (O.Z.), the Canadian Natural Sciences and Engineering Research Council (NSERC) award RGPIN 2018-03898 (J.T.B.), NSERC award RGPIN-2017-04636 (A.S.L.). S.K. was partially supported by funding from the Memorial University of Newfoundland School of Graduate Studies.

The views and conclusions contained in this document are those of the authors and should not be interpreted as necessarily representing the official policies, either expressed or implied, of the U.S. Department of Health and Human Services or of the institutions and companies affiliated with the authors, nor does mention of trade names, commercial products, or organizations imply endorsement by the U.S. Government.

