## Supplementary Tables S1-S6 for "Formal Recognition and Classification of Gene Transfer Agents as Viriforms"

for

**Formal Recognition and Classification of Gene Transfer Agents as Viriforms**

*Roman Kogay[1], Sonja Koppenhöfer[2], J. Thomas Beatty[3], Jens H. Kuhn[4], Andrew S. Lang[2], and Olga Zhaxybayeva[1,#]*

*[1]Department of Biological Sciences, Dartmouth College, Hanover, NH, USA; [2]Department of Biology, Memorial University of Newfoundland, St. John's, NL, Canada; [3]Department of Microbiology & Immunology, University of British Columbia, Vancouver, BC, Canada; [4]Integrated Research Facility at Fort Detrick, Division of Clinical Research, National Institute of Allergy and Infectious Diseases, National Institutes of Health, Frederick, MD, USA*

**Supplementary Table S1. Rhodobacter capsulatus gene transfer agent (RcGTA) genes, their locus tags, and their functional annotations in the host (*Rhodobacter capsulatus* SB1003) genome (GenBank accession CP001312.1).**

| Locus tag | Start | End | Strand | GenBank annotation* |
|---|---|---|---|---|
| RCAP_rcc00171 | 203033 | 204148 | + | tail fiber protein [1] |
| RCAP_rcc00555 | 593866 | 594477 | + | holin-associated N-acetylmuramidase |
| RCAP_rcc00556 | 594468 | 594962 | + | holin family protein |
| RCAP_rcc01079 | 1149611 | 1149865 | + | *ghsA* head spike protein [2] |
| RCAP_rcc01080 | 1149862 | 1150839 | + | *ghsB* head spike protein [2] |
| RCAP_rcc01682 | 1818689 | 1819012 | + | small terminase [3] |
| RCAP_rcc01683 | 1818963 | 1820309 | + | terminase family protein |
| RCAP_rcc01684 | 1820495 | 1821685 | + | phage portal protein |
| RCAP_rcc01685 | 1821695 | 1821922 | + | hypothetical protein |
| RCAP_rcc01686 | 1821933 | 1822487 | + | HK97 family phage prohead protease |
| RCAP_rcc01687 | 1822519 | 1823715 | + | phage major capsid protein |
| RCAP_rcc01688 | 1823888 | 1824481 | + | adaptor protein [4] |
| RCAP_rcc01689 | 1824478 | 1824816 | + | head-tail adaptor protein |
| RCAP_rcc01690 | 1824813 | 1825220 | + | tail terminator protein [4] |
| RCAP_rcc01691 | 1825261 | 1825674 | + | phage major tail protein, TP901-1 family |
| RCAP_rcc01692 | 1825682 | 1826008 | + | gene transfer agent family protein |
| RCAP_rcc01693 | 1826005 | 1826232 | + | phage tail assembly chaperone |
| RCAP_rcc01694 | 1826219 | 1826878 | + | phage tail tape measure protein |
| RCAP_rcc01695 | 1826889 | 1827521 | + | distal tail protein [4] |
| RCAP_rcc01696 | 1827521 | 1828411 | + | baseplate hub protein [4] |
| RCAP_rcc01697 | 1828408 | 1828860 | + | peptidase |
| RCAP_rcc01698 | 1828861 | 1832775 | + | glycoside hydrolase/phage tail family protein |
| RCAP_rcc01865 | 2018987 | 2020135 | - | *gafA* transcriptional regulator [5] |
| RCAP_rcc01866 | 2020352 | 2020660 | + | Possibly, maturation/assembly protein [1] |

* For some genes, the GenBank annotations in the host genome record were replaced with more informative functional annotations from experimental work, with relevant citations provided.

**Supplementary Table S2. Dinoroseobacter shibae gene transfer agent (DsGTA) genes, their locus tags, and their functional annotations in the host (*Dinoroseobacter shibae* DFL12) genome (GenBank accession CP000830.1).**

| Locus tag | Start | End | Strand | GenBank annotation |
|---|---|---|---|---|
| Dshi_1584 | 1649803 | 1650927 | - | DUF6456 domain-containing protein |
| Dshi_1585 | 1651121 | 1651423 | + | DUF6477 family protein |
| Dshi_1756 | 1820645 | 1821166 | - | holin family protein |
| Dshi_1757 | 1821157 | 1821768 | - | holin-associated N-acetylmuramidase |
| Dshi_2162 | 2291686 | 2292393 | - | DUF2793 domain-containing protein |
| Dshi_2163 | 2292386 | 2296303 | - | glycoside hydrolase/phage tail family protein |
| Dshi_2164 | 2296307 | 2296747 | - | peptidase |
| Dshi_2165 | 2296744 | 2297634 | - | DUF2163 domain-containing protein |
| Dshi_2166 | 2297631 | 2298263 | - | DUF2460 domain-containing protein |
| Dshi_2167 | 2298277 | 2298945 | - | phage tail tape measure protein |
| Dshi_2168 | 2298929 | 2299144 | - | phage tail assembly chaperone |
| Dshi_2169 | 2299141 | 2299470 | - | gene transfer agent family protein |
| Dshi_2170 | 2299476 | 2299889 | - | phage major tail protein, TP901-1 family |
| Dshi_2171 | 2299911 | 2300321 | - | DUF3168 domain-containing protein |
| Dshi_2172 | 2300318 | 2300656 | - | head-tail adaptor protein |
| Dshi_2173 | 2300653 | 2301243 | - | hypothetical protein |
| Dshi_2174 | 2301398 | 2302615 | - | phage major capsid protein |
| Dshi_2175 | 2302612 | 2303229 | - | HK97 family phage prohead protease |
| Dshi_2176 | 2303242 | 2303502 | - | hypothetical protein |
| Dshi_2177 | 2303499 | 2304689 | - | phage portal protein |
| Dshi_2178 | 2304797 | 2306161 | - | terminase family protein |

**Supplementary Table S3. Rhodovulum sulfidophilum gene transfer agent (RsGTA) genes, their locus tags, and their functional annotations in the host (*Rhodovulum sulfidophilum* DSM 1374) genome (GenBank accession CP015418.1).**

| Locus tag | Start | End | Strand | GenBank annotation |
|---|---|---|---|---|
| A6W98_RS21760 | 181175 | 182050 | + | DUF2793 domain-containing protein |
| A6W98_09125 | 1925273 | 1925629 | + | hypothetical protein |
| A6W98_09130 | 1925622 | 1926899 | + | terminase family protein |
| A6W98_09135 | 1927031 | 1928206 | + | phage portal protein |
| A6W98_09140 | 1928209 | 1928451 | + | hypothetical protein |
| A6W98_09145 | 1928472 | 1929038 | + | HK97 family phage prohead protease |
| A6W98_09150 | 1929082 | 1930269 | + | phage major capsid protein |
| A6W98_09155 | 1930432 | 1931031 | + | hypothetical protein |
| A6W98_09160 | 1931028 | 1931369 | + | head-tail adaptor protein |
| A6W98_09165 | 1931366 | 1931776 | + | DUF3168 domain-containing protein |
| A6W98_09170 | 1931800 | 1932213 | + | phage major tail protein, TP901-1 family |
| A6W98_09175 | 1932216 | 1932548 | + | gene transfer agent family protein |
| A6W98_09180 | 1932538 | 1932732 | + | phage tail assembly chaperone |
| A6W98_09185 | 1932736 | 1933404 | + | phage tail tape measure protein |
| A6W98_09190 | 1933420 | 1934052 | + | DUF2460 domain-containing protein |
| A6W98_09195 | 1934052 | 1934939 | + | DUF2163 domain-containing protein |
| A6W98_09200 | 1934936 | 1935379 | + | NlpC/P60 family protein |
| A6W98_09205 | 1935383 | 1939282 | + | glycoside hydrolase/phage tail family protein |
| A6W98_10845 | 2293061 | 2293672 | + | holin-associated N-acetylmuramidase |
| A6W98_10850 | 2293663 | 2294229 | + | holin family protein |
| A6W98_12585 | 2663257 | 2663559 | - | DUF6477 family protein |
| A6W98_12590 | 2663793 | 2664887 | + | DUF6456 domain-containing protein |

**Supplementary Table S4. Ruegeria pomeroyi gene transfer agent (RpGTA) genes, their locus tags, and their functional annotations in the host (*Ruegeria pomeroyi* DSS-3) genome (GenBank accession CP000031.2).**

| Locus tag | Start | End | Strand | GenBank annotation |
|---|---|---|---|---|
| SPO1878 | 2001541 | 2002152 | + | holin-associated N-acetylmuramidase |
| SPO_RS09550 | 2002143 | 2002721 | + | holin family protein |
| SPO2106 | 2242168 | 2243322 | - | DUF6456 domain-containing protein |
| SPO2107 | 2243566 | 2243874 | + | DUF6477 family protein |
| SPO2250 | 2393488 | 2397414 | - | glycoside hydrolase/phage tail family protein |
| SPO2251 | 2397414 | 2397860 | - | peptidase |
| SPO2252 | 2397857 | 2398744 | - | DUF2163 domain-containing protein |
| SPO2253 | 2398744 | 2399376 | - | DUF2460 domain-containing protein |
| SPO2254 | 2399392 | 2400054 | - | phage tail tape measure protein |
| SPO2255 | 2400041 | 2400250 | - | phage tail assembly chaperone |
| SPO2256 | 2400247 | 2400567 | - | gene transfer agent family protein |
| SPO2257 | 2400570 | 2400983 | - | phage major tail protein, TP901-1 family |
| SPO2258 | 2401008 | 2401421 | - | DUF3168 domain-containing protein |
| SPO2259 | 2401418 | 2401756 | - | phage head closure protein |
| SPO2260 | 2401753 | 2402352 | - | head-tail connector protein |
| SPO2261 | 2402517 | 2403695 | - | phage major capsid protein |
| SPO2262 | 2403729 | 2404286 | - | HK97 family phage prohead protease |
| SPO2263 | 2404315 | 2404539 | - | hypothetical protein |
| SPO2264 | 2404532 | 2405704 | - | phage portal protein |
| SPO2266 | 2405830 | 2407056 | - | terminase family protein |
| SPO_RS11495 | 2407109 | 2407453 | - | hypothetical protein |
| SPO2726 | 2911999 | 2913924 | - | DUF2793 domain-containing protein |

**Supplementary Table S5. Bartonella gene transfer agent (BaGTA) genes, their locus tags, and their functional annotations in the host (*Bartonella henselae* str. Houston-1) genome (GenBank accession BX897699.1).**

| Locus tag | Start | End | Strand | GenBank annotation* |
|---|---|---|---|---|
| BH13940 | 1602076 | 1602738 | - | phage related lysozyme |
| BH13950 | 1603017 | 1603283 | - | hypothetical protein |
| BH13960 | 1604763 | 1605770 | - | phage tail collar protein [1] |
| BH13970 | 1605767 | 1606891 | - | peptidase, chaperon [1] |
| BH13990 | 1607638 | 1609101 | - | hypothetical protein |
| BH14000 | 1610017 | 1610712 | - | hypothetical protein |
| BH14010 | 1611092 | 1612210 | - | major capsid protein [1] |
| BH14040 | 1613196 | 1613993 | - | hypothetical protein |
| BH14060 | 1614966 | 1616819 | - | hypothetical protein |
| BH14070 | 1616804 | 1618012 | - | phage terminase [1] |
| BH14090 | 1618545 | 1618874 | - | hypothetical protein |
| BH14150 | 1625365 | 1626306 | + | hypothetical protein |
| BH14200 | 1630937 | 1631227 | + | hypothetical protein |
| BH14220 | 1631403 | 1632176 | + | hypothetical protein |
| BH14240 | 1632865 | 1633209 | - | hypothetical protein |
| BH14310 | 1638603 | 1638968 | + | hypothetical protein |
| BH14320 | 1639193 | 1639567 | + | hypothetical protein |
| BH14330 | 1639687 | 1639965 | + | hypothetical protein |
| BH14470 | 1655075 | 1655464 | - | hypothetical protein |
| BH14480 | 1655778 | 1656095 | - | hypothetical protein |
| BH14490 | 1656650 | 1657579 | + | hypothetical protein |
| BH14500 | 1657576 | 1657869 | + | hypothetical protein |
| BH14510 | 1657853 | 1658998 | + | hypothetical protein |
| BH14520 | 1658982 | 1661741 | + | phage related protein |
| BH14530 | 1663283 | 1663849 | + | hypothetical protein |
| BH14580 | 1676092 | 1676691 | + | hypothetical protein |
| BH14600 | 1679632 | 1680294 | + | phage related lysozyme |
| BH14610 | 1680291 | 1680518 | + | hypothetical protein |
| BH14620 | 1680647 | 1680868 | + | hypothetical protein |

* For some genes, the GenBank annotations in the host genome record were replaced with more informative functional annotations from experimental work, with relevant citations provided.

1. Quebatte, M., Christen, M., Harms, A., Korner, J., Christen, B., and Dehio, C. Gene transfer agent promotes evolvability within the fittest subpopulation of a bacterial pathogen. *Cell Syst*, 2017. 4(6): 611-621.e6.

**Supplementary Table S6. Brachyspira hyodysenteriae gene transfer agent (BhGTA) genes, their locus tags, and their functional annotations in the host (*Brachyspira hyodysenteriae* WA1) genome (GenBank accession CP001357.1).**

| Locus tag | Start | End | Strand | GenBank annotation* |
|---|---|---|---|---|
| BHWA1_01838 | 2121760 | 2123001 | + | hypothetical protein |
| BHWA1_01839 | 2123304 | 2124257 | + | hypothetical protein |
| BHWA1_01840 | 2124448 | 2124816 | + | hypothetical protein |
| BHWA1_01841 | 2125122 | 2126165 | + | major capsid protein [1] |
| BHWA1_01844 | 2127442 | 2127999 | + | hypothetical protein |
| BHWA1_01846 | 2128455 | 2129852 | + | hypothetical protein |
| BHWA1_01850 | 2131660 | 2132535 | + | hypothetical protein |
| BHWA1_01851 | 2132609 | 2135149 | + | hypothetical protein |
| BHWA1_01852 | 2135161 | 2135709 | + | hypothetical protein |
| BHWA1_01853 | 2135714 | 2136307 | + | endolysin [1] |
| BHWA1_01855 | 2137181 | 2137438 | + | holin [1] |

* For some genes, the GenBank annotations in the host genome record were replaced with more informative functional annotations from experimental work, with relevant citations provided.

1. Matson, E.G., Thompson, M.G., Humphrey, S.B., Zuerner, R.L., and Stanton, T.B. Identification of genes of VSH-1, a prophage-like gene transfer agent of *Brachyspira hyodysenteriae*. *J Bacteriol*, 2005. **187**(17): 5885-92.
